# The origins and diversification of Holarctic brown bear populations inferred from genomes of past and present populations

**DOI:** 10.1101/2023.02.08.527755

**Authors:** Takahiro Segawa, Alba Rey-Iglesia, Eline D Lorenzen, Michael V Westbury

## Abstract

The brown bear (*Ursus arctos*) is one of the survivors of the Late Quaternary megafauna extinctions. However, despite being widely distributed across the Holarctic, brown bears have experienced extensive range reductions, and even extirpations in some geographic regions. Previous research efforts utilising genetic data have provided valuable insights into their evolutionary history. However, most studies have been limited to contemporary individuals or mitochondrial DNA, limiting insights into population processes that preceded the present. Here we present genomic data from two Late Pleistocene brown bears from Honshu, Japan, and eastern Siberia, and combine them with published contemporary and ancient genomes from across the Holarctic range of brown bears to investigate the evolutionary relationships among brown bear populations through time and space. By including genomic data from Late Pleistocene and Holocene individuals sampled outside the current distribution range we uncover diversity not present in the contemporary populations. Notably, although contemporary individuals display geographically structured populations most likely driven by isolation-by-distance, this pattern varies among the ancient samples across different regions. The inclusion of ancient brown bears in our analysis provides novel insights into the evolutionary history of brown bears and contributes to understanding the populations and diversity lost during the Late Quaternary.

## Introduction

The Pleistocene-Holocene transition was a period of major climatic upheaval, culminating with a large number of megafaunal extinctions as a defining feature across the globe [1]. However, despite this, many megafauna species still persist today, although they may have experienced local extirpations, range reductions, and losses of genetic diversity [2,3]. By comparing the genetic composition of populations inhabiting recently recolonised regions to that of populations located in past refugia, genetic information from contemporary individuals can be used to study the patterns of postglacial recolonisation after the Last Glacial Maximum [4]. However, purely focussing on contemporary individuals means results are restricted to surviving lineages, limiting our understanding of the population processes that preceded the present. Genetic information from subfossil individuals (ancient DNA - aDNA) enables the tracking of change in real time, the uncovering of potentially lost lineages, and an overall better understanding of past responses to change [5].

The brown bear (*Ursus arctos*) is one of the survivors of the Late Pleistocene megafauna extinctions. The species is one of the largest extant terrestrial carnivores, and is distributed across the Holarctic [6]. The ability of brown bears to survive may be due to their ecological flexibility and broad dietary range, observed in contemporary populations [7]. Brown bears have evolved a generalist omnivore strategy; although they possess morphological traits characteristic of carnivores, their diets in many ecosystems comprise primarily of plant matter [8]. Despite their current dietary flexibility, brown bears experienced extensive range reductions during the Holocene, with extirpations in some geographic regions driven by climate and environmental change, and more recently, humans [9]. For example, based on fossil chronology, the relatively large steppe brown bear (*Ursus arctos priscus*) of Eurasia disappeared ∼25 kya. Although it was presumably well-adapted for survival in Europe during the Late Pleistocene, its size may have inhibited adaptive flexibility after the environment drastically changed at the onset of the Holocene [10].

Since its first utilisation in brown bears in 1994 [11], genetics has provided many insights into the contemporary phylogeographic structuring of the species. However, the majority of mitochondrial genetic clades used to form biogeographic hypotheses are paraphyletic, and not unique to a single geographic region [12], making global inferences difficult. A recent study using nuclear genomes from across the contemporary Holarctic distribution showed phylogeographic patterns at the nuclear level do not mirror the mitochondria, but instead found geographically structured populations following isolation by distance. They suggested that current population structure may represent recent population dynamics [13], as more ancient signals may have been broken down by gene flow and recombination.

Ancient DNA from brown bears has provided valuable insights into their evolutionary history, and provided valuable information regarding temporal changes in genetic diversity and migration patterns [14–17]. However, studies of ancient brown bear specimens have thus far been based on mitochondrial DNA, which represents just a single loci, is exclusively maternally inherited, and may be heavily confounded by sex-biased migrations, gene flow, and incomplete lineage sorting [13,18,19]. Nuclear data provides an opportunity to circumvent the challenges associated with mitochondrial genomes [18]. Although ancient nuclear genomic data is available for several brown bears, it has thus far only been used to understand the evolutionary relationships between different Ursine bear species [20,21], not to understand brown bear intraspecific relationships through time.

Here, we utilise a combination of contemporary and ancient nuclear genomes, incorporating newly generated palaeogenomic data from two Late Pleistocene individuals from Japan (0.11x; 32,464 ± 255 cal. BP [14]) and eastern Siberia (0.35x; molecularly dated to 61,826 years BP [HPD 95%: 27,866–96,783 years BP][22]), to investigate the evolutionary relationships among brown bear populations through time and space.

## Methods

### Data generation

We generated palaeogenomic data from a Late Pleistocene (32,464 ± 255 cal. BP [14]) brown bear from the Japanese island of Honshu (JBB-32K [14]). Lab work was carried out in a clean room for ancient DNA at the National Institute of Polar Research, Japan. We extracted 100 mg of petrosal bone, and followed the DNA extraction procedure based on silica pellets in solution as described in Orlando et al. [23]. Double-stranded DNA was converted into an Illumina sequencing library using an NEBNext Quick DNA Library Prep Master Mix Set for 454 (New England BioLabs) with 20 μl of DNA extract without DNA fragmentation. Illumina sequencing adapters were added to the end-repair reaction. Indexing PCR was prepared in the ancient DNA laboratory, carried out in a physically separate modern laboratory. Each eluate (3 µl) was subjected to PCR amplification in 50-μl reactions containing 1 KAPA HiFi HotStart Uracil+ReadyMix (Kapa Biosystems) and 0.3 μM Dual Index primers of NEBNext Multiplex Oligos for Illumina (NEB). Three independent PCR reactions from the same library were combined after PCR and purified using a NucleoSpin Gel and PCR Clean-up kit (Macherey-Nagel). Sequencing was carried out on an Illumina HiSeq XTen at Macrogen, Japan. Previously published data for this individual was downloaded from DDBJ DRA IDs: DRR308533 and DRR308534 [14].

We generated palaeogenomic data from a Late Pleistocene (aged beyond radiocarbon dating: >50 kya but molecularly dated to 61,826 years BP [HPD 95%: 27,866–96,783 years BP][22]) brown bear from eastern Siberia (CGG_1_020006 [22]). Lab work was carried out in a clean room for ancient DNA at Globe Institute, University of Copenhagen. We extracted DNA from ∼50 mg of bone powder using a modified version of the protocol from [24]. Modifications included overnight incubation with the extraction buffer at 42□°C instead of at 37□°C, the bone powder was pelleted out of suspension, and the supernatant concentrated down to 150–200 μl for each sample using 30 kDa Amicon centrifugal filter unit (Millipore). The remaining supernatant was mixed with 13x its volume of binding buffer and DNA was purified with MinElute columns (Qiagen), following the manufacturer’s instructions with the exception of a 15-minute incubation at 37□°C during the elution step. DNA extracts were transformed into sequencing libraries in 25 μl reactions following the protocol from Meyer and Kircher [25] with the modification that the initial DNA fragmentation was not performed, as ancient DNA is already highly fragmented, and MinElute kit (Qiagen) was used for the purification steps. DNA libraries were indexed using KAPA HiFi uracil[+[premix (KAPA Biosystems). The number of cycles for index PCRs was determined from qPCR analysis. The resulting libraries were quantified on an Agilent 2100 Bioanalyzer, pooled at equimolar concentration and sequenced on an Illumina HiSeq 2500 at the University of Copenhagen using single-end 80 base pairs (bp) reads.

We additionally downloaded the raw reads from 137 contemporary and eight ancient brown bear individuals, from across their present and past distributions [13,20,21,26–31] (Fig 1A and Supplementary Table 1). Seven of the ancient samples are from outside the current distribution of brown bears, and include Ireland (n = 5), Honshu (Japan, n = 1), and Quebec (Canada, n = 1).

**Figure 1:**
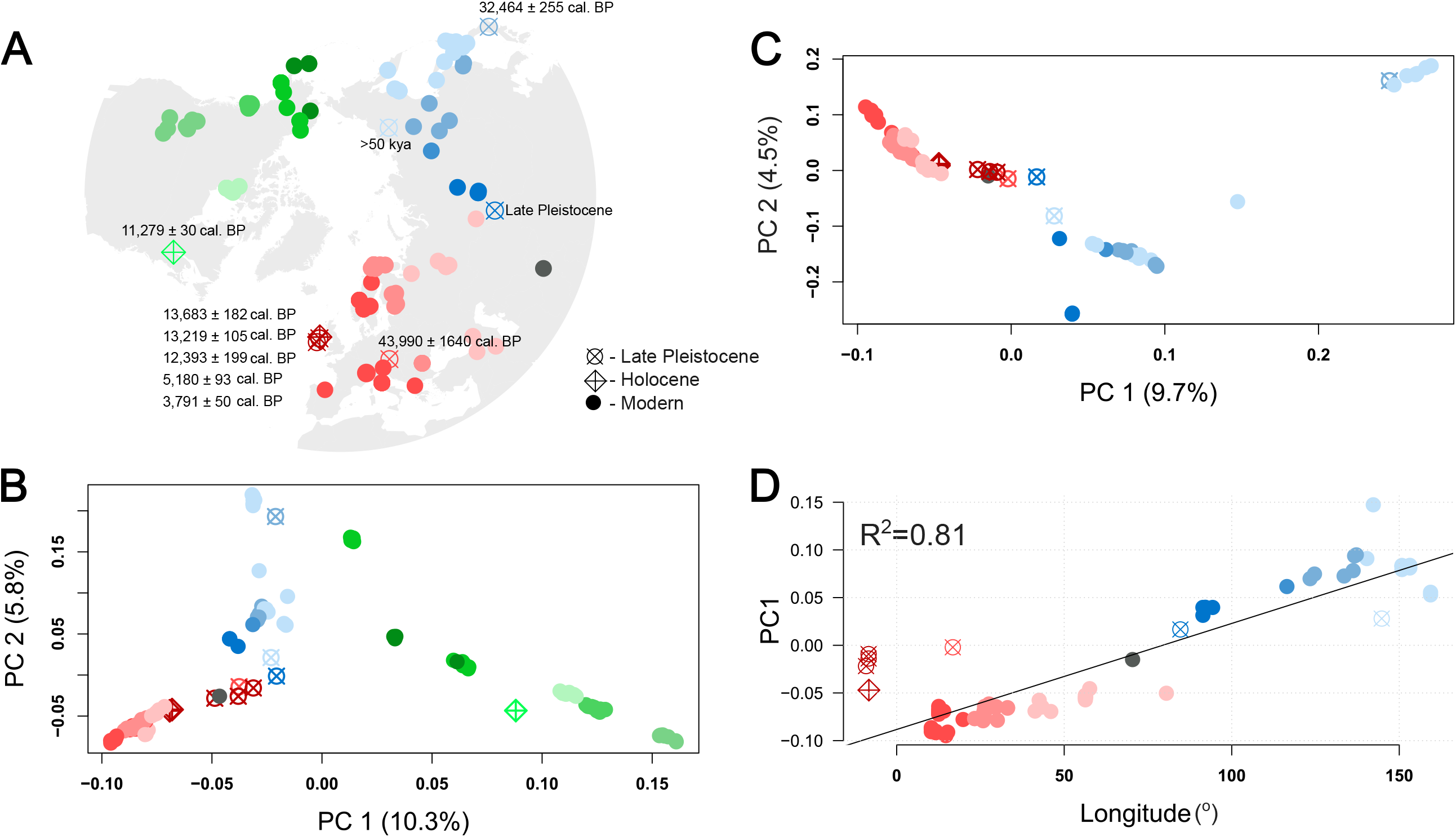
Brown bear sample localities and population structure. A) Map showing the localities of the brown bear ancient and contemporary brown bear genomes analysed. Calibrated radiocarbon ages of the ancient samples included are listed next to their locality. ‘Late Pleistocene’ next to Denisova_LP indicates a direct age is unknown, but it is listed as Late Pleistocene in the original publication [26]. CGG_1_020006 was dated to beyond the limit of radiocarbon dating (>50 kya) but molecularly dated to 61,826 years BP [HPD 95%: 27,866–96,783 years BP][23]. B) PCA generated from genotype likelihoods of all samples. C) PCA generated from genotype likelihoods of the Eurasian samples only. Percentage values in parentheses on the X and Y axes show the percentage of variation explained by the principal component. Colours and symbols represent geography and age, respectively. D) PC1 vs degrees longitude for the Eurasian individuals, excluding the Japanese individuals. Dark grey circle shows the single Himalayan individual. The colour of CentralRussia1 was changed to light red due to its geographic position in the Ural mountains and European-like clustering in the PCA.

### Data processing and mapping

#### Contemporary brown bears

We processed the raw sequencing data from 35 contemporary brown bears (excluding those from Endo et al and de Jong et al, detailed below) with the PALEOMIX v.1.2.12 [32] pipeline. Adapters, stretches of Ns, and low-quality bases in reads were trimmed and filtered with AdapterRemovalv2 [33] using default parameters. BWA-backtrack v0.7.15 [34] was used to map the cleaned reads to the polar bear pseudo-chromosome genome [35,36] (based on Genbank accession: GCA_000687225.1), which includes a mitochondrial genome (Genbank accession: NC003428), with default parameters. Reads with mapping quality < 30 were filtered using SAMtools [37]. Duplicates were removed with Picard v2.6.0 [38]. Possible paralogs were filtered using SAMtools v1.6. Finally, local realignment around indels was done using GATK (v 3.3) [39].

The individuals from Endo et al (n=6) and de Jong et al (n=96) followed a similar protocol, but using Fastp v0.23.2 [40] for adapter trimming and duplicate removal using SAMtools. To be used as an outgroup/ancestral sequence, the raw data for the spectacled bear (*Tremarctos ornatus*) was downloaded (SRA accession code: SRR13602719) and mapped to the same polar bear reference genome as above using the BWA mem algorithm.

#### Ancient brown bears

We removed illumina adapter sequences, low-quality reads (mean q <25), short reads (<30bp), and merged overlapping read pairs with Fastp. We mapped the merged reads to the polar bear reference genome using BWA-backtrack v0.7.15, with the seed disabled (-l 999), and otherwise default parameters. Reads with mapping quality < 30 and duplicates were filtered using SAMtools [37].

#### Population structure

We evaluated the relationships between the contemporary and ancient brown bears included in this study by performing Principal Component Analyses (PCA). We performed these using either pseudohaploid base calls or genotype likelihoods computed with ANGSDv0.921 [41]. We computed genotype likelihoods in ANGSD for all individuals specifying the parameters: minimum mapping and base qualities of 30 (-minmapQ 30 -minQ 30), calculate genotype likelihoods using the GATK algorithm (-GL 2), output a beagle genotype likelihood file (-doGlf 2), calculate major and minor alleles based on genotype likelihoods (-doMajorMinor 1), remove transitions (-rmtrans 1), only include SNPs with a p-values <1e-6 (-SNP_pval 1e-6), only consider autosomal chromosomes (-rf), a minimum minor allele frequency of 0.05 (-minmaf 0.05), skip triallelic sites (-skiptriallelic 1), only consider reads mapping to one region uniquely (-uniqueonly 1), and only consider sites if at least 132 individuals have coverage (-minind 132). We only included pseudochromosomes 1-18 as previously recommended [35]. The pseudohaploid base call was computed using the same parameters with the addition of -doIBS 2, -doCov 1, and only included SNPs if at least two individuals had the minor allele (-minminor 2). To construct covariance matrices from the genotype likelihoods datasets we used PCAngsd v0.98 [42]. We only included a single Apennines mountain (Italy) brown bear in these analyses due to their high levels of relatedness [29]. We repeated these steps with only individuals from Eurasia using the same parameters but set the minimum individual parameter (MinInd) to 79. Due to its suitability for high levels of missing data, we additionally performed principal component analyses for the two datasets using EMU v0.9 [43]. As input, we generated consensus pseudohaploid base calls in ANGSD using the same parameters as above but with the addition of the –dohaplocall 2. We converted the haploid calls into PLINK format using haploToPlink, which is part of the ANGSD toolkit. We ran the PLINK file in EMU specifying to produce eigenvalues for the first three principal components, and otherwise default parameters.

To investigate whether the low coverage, and ancient DNA damage may influence the ancient individual’s placement within a PCA, we simulated ancient DNA damage patterns, short read lengths, and low coverage onto three contemporary individuals. The three individuals were S235 (Central Asia) and Kirkenes (Europe), and DRR276778 (Japan). We used Fastp to reduce the raw reads to 40bp then added ancient DNA patterns to the raw reads of these three individuals using TAPAS v1.2 [44]. We mapped the artificially damaged and short reads to the reference genome following the approaches specified above and we used SAMtools to subsample the individuals. Ancient DNA damage patterns were taken from the Sligo-14 mapdamage output for S235 and Kirkenes, and JBB-32K for DRR276778. We downsampled S235 and Kirkenes to 0.1x and JBB-32K to 0.05x. We reran the genotype likelihoods and pseudohaploid PCA with these three individuals replacing the original high-quality data with the artificially damaged bam files.

#### Pairwise distances

To further estimate the relationships between ancient and contemporary Eurasian individuals, we calculated pairwise distances between all Eurasian individuals (excluding JBB-32K due to its clear placement within the Hokkaido brown bears in the PCA) using the pseudohaploid baseball (-doIBS 2) mentioned above and converted these into a distance matrix using the -makematrix 1 parameter in ANGSD. We extracted the columns corresponding to the ancient individuals and rows corresponding to contemporary individuals.

#### Ancestry proportions

We calculated the ancestry proportions of the eight ancient Eurasian individuals (excluding JBB-32K) using admixfrog v0.7.2 [45]. For this analysis we considered three ancestry states each represented by a single individual; East Europe - represented by Kirkenes, Himalayas - represented by Himalaya1, and Central Asia - represented by S235. As input for the reference panel we created a multi-individual variant call file with the three individuals using BCFtools v1.15 [46], specifying chromosomes 1-18 (-f), minimum mapping and base qualities of 20 (-q 20 -Q 20). We ran admixfrog specifying a minimum read length of 30bp, a bin size of 50kb, and otherwise default parameters. When interpreting the results, if a window was returned as having combined ancestry between any of the two states, then we deemed this as ancestral.

#### D-statistics topology test

We used D-statistics to investigate whether our ancient Eurasian individuals were more closely related to contemporary European or Asian individuals. D-statistics works by calculating shared derived alleles between non-sister branches of a given topology [[[H1, H2], H3], Outgroup]. An allele is considered as derived if it is different to the outgroup allele. Although it is most commonly implemented to investigate differential levels of gene flow between non-sister taxon, it can also be used to investigate shared polymorphisms and population structure [47]. We subsampled the dataset to 42 Eurasian individuals (Supplementary table S2). We calculated the D-statistics using a random base call approach in ANGSD (-doabbababa 1), specifying only pseudochromosomes 1-18, the spectacled bear as outgroup, and the following parameters: -minmapQ 30 -minQ 30 -blocksize 1000000 - rmtrans 1 -uniqueonly 1. We summarised the results using a block jackknife approach with the Rscript available in the ANGSD toolsuite (ANGSD_jackknife.R). As ANGSD computes every possible ingroup triplet combination of D-statistics, we filtered out only the combinations of interest. We filtered the output to only include the contemporary European individuals in the H1 position, the contemporary Asian individuals in H2, and the ancient individuals in the H3 position [[[Europe, Asia], Ancient], Outgroup]. This filtering allows us to see if the ancient individual has more shared derived alleles with either the contemporary European or contemporary Asian individual.

## Results

### Population structure

Regardless of the method used (genotype likelihood, pseudohaploid, EMU) we found similar results in our PCAs, with clear differences between bears from different geographic localities in a west-to-east cline (Fig 1B,C, Supplementary Fig S1). Further investigations into Eurasia showed a very strong correlation between PC1 and longitude (R^2^=0.81, Fig 1D). The ∼32kya Honshu Japanese individual (JBB-32K) clustered most closely with the contemporary Hokkaido Japanese bears. The ∼11kya Quebec sample clustered with the North American bears, but not with any specific locality in the genotype likelihoods or pseudohaploid data. However, it did cluster with the easternmost bears in the EMU-computed PCA.

The remaining eight ancient individuals did not cluster where expected based on geography. The Holocene Irish individuals (Leitrim-4, and Leitrim-5) clustered close to the easternmost European individuals, whereas the Late Pleistocene Irish (Clare-12, Sligo-13, Sligo-14), Austrian (Austria_LP), and Siberian (Denisova_LP, CGG_1_020006) cluster closest to the Himalayan individual (seen as dark grey in Fig 1A) but in general do not cluster within any contemporary individual.

Investigations into the role of ancient DNA damage in the placement of individuals into the PCA showed that simulated ancient individuals clustered in the same location as they did when using the high-quality version of the data, and that using genotype likelihoods may be the most robust to aDNA damage (Supplementary Fig S2). This suggests that our approaches are robust to aDNA damage and the results are biological rather than driven by data bias.

When visualising PC3 in the Eurasian dataset, the differentiation between contemporary European individuals became more apparent, yet retained the structuring observed in PC1.

### Pairwise distances

Overall, the Irish bears generally show the lowest pairwise distances (PWD) to individuals from Europe, but this heavily overlaps with some Asian bears (Fig 2A). Austria_LP and Denisova_LP show the smallest PWD to the Himalayan individual. However, when excluding the Himalayas bear, Austria_LP is more or less similar to all bears, and Denisova_LP is closer to Asian bears than European. PWD heavily overlapped between CGG_1_020006 and Europe, Asia, and the Himalayas.

**Fig 2.**
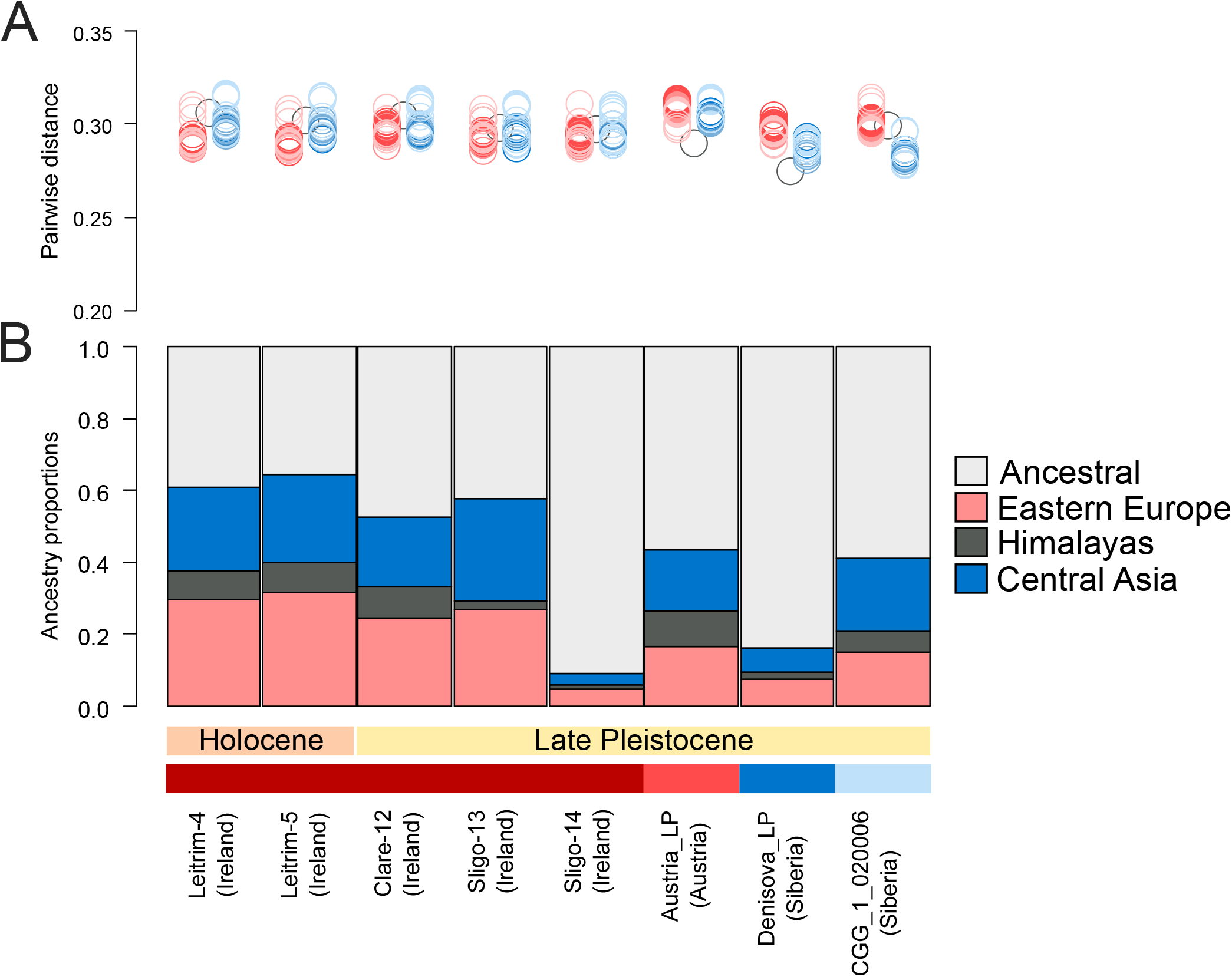
Relationship of ancient Eurasian individuals to contemporary individuals. A) Genome-wide pairwise distances to contemporary individuals. B) Ancestry proportions calculated using three reference populations. Ancestral is defined as having combined ancestry between any of the two reference populations. Coloured boxes above the names on the X axis correspond to the geographic location (longitude) of the ancient sample as seen in Fig 1A.

### Ancestry proportions

We found relatively high proportions of most of the ancient Irish individual’s ancestry (to the exclusion of Sligo-14) to be eastern European, followed by central Asia, then Himalayan (Fig 2B). There was slightly more European ancestry in the Holocene Irish individuals (Leitrim-4 0.297 and Leitrim-5 0.318) relative to their Late Pleistocene counterparts (Clare-12 0.246, Sligo-13 0.27, Sligo-14 0.046). The lowest level of ancestry in all ancient individuals - when considering only the three single reference populations - was Himalayan (<0.09). Large proportions of ancestry in all ancient individuals could not be assigned to a single population, especially in Sligo-14 (0.91) and Denisova_LP (0.837), so was denoted as ancestral. Austria_LP had similar values of eastern European and central Asian ancestry (0.165 and 0.168 respectively), whereas CGG_1_020006 had slightly more central Asian (0.205) relative to eastern European or Himalayan (0.149 and 0.059 respectively).

### D-statistics topology test

All ancient Irish individuals, regardless of age, show similar D-statistics results (Fig 3). However, there is a slight shift towards more negative D-scores in the Holocene individuals relative to the LP individuals, indicating more European ancestry in the Holocene individuals. The mean D-scores of the Late Pleistocene individuals was -0.050, -0.056, -0.052 for Clare-12, Sligo-13, and Sligo-14 respectively and the mean D-scores of the Holocene individuals was -0.077, -0.076 for Leitrim-4 and Leitrim-5, respectively. The ancient Asian individuals, CGG_1_020006 and JBB-32K, showed more positive values when including Asian individuals in the H2 position, suggesting a closer relationship to Asia than to Europe (Fig 3). In contrast, Denisova_LP showed a clustering of values around zero, suggesting no obvious differential affinity to either contemporary European or Asian individuals. However, the mean D-score of 0.01 may indicate a slightly higher affinity to contemporary Asian individuals. The ancient Austria_LP individual also showed similar D-scores between H2 individuals, with a clustering of values close to zero, but an overall tendency towards negative D-scores (mean D-score -0.01) suggesting more European ancestry (Fig 3).

**Fig 3.**
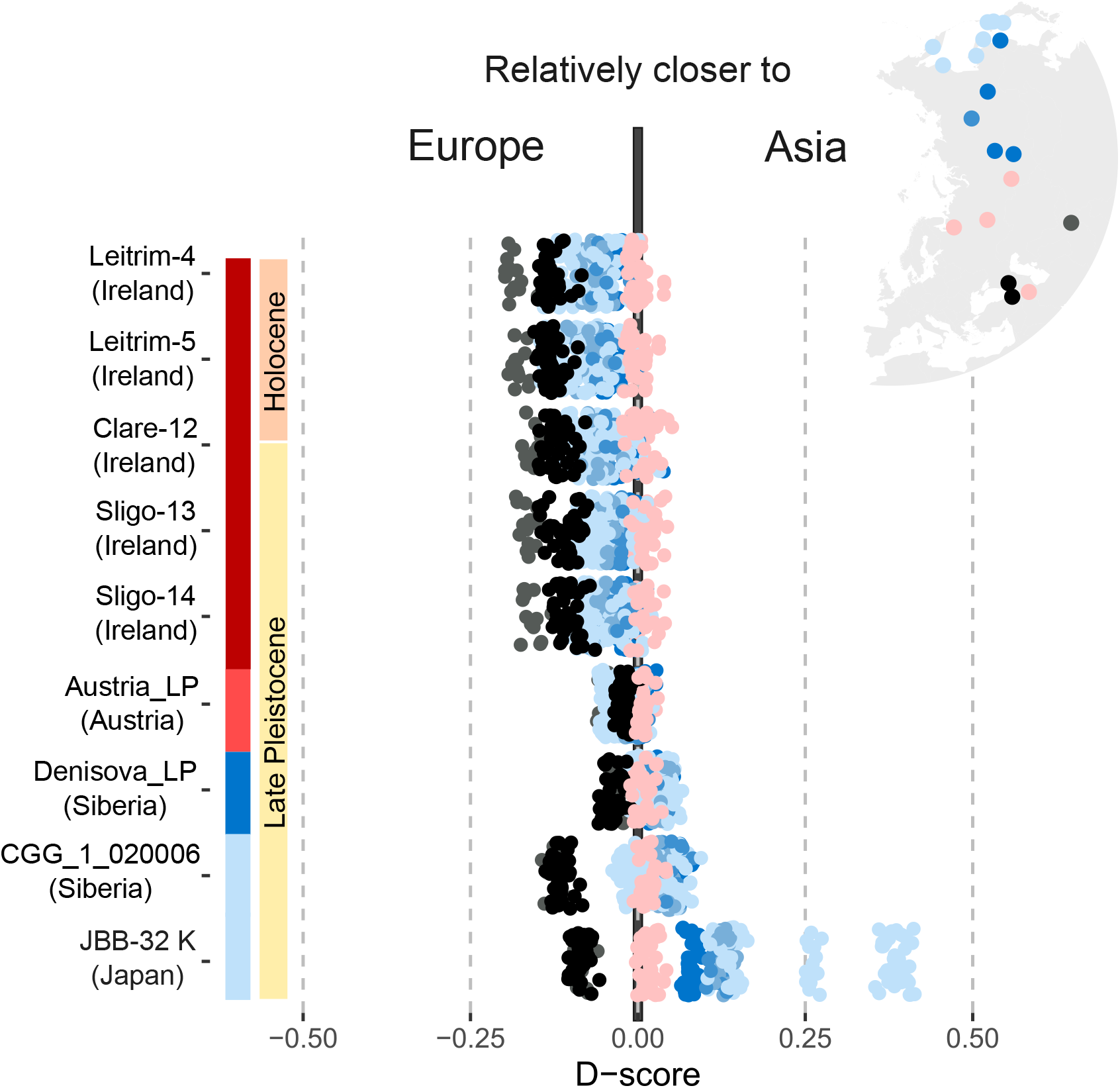
Test for population structure using D-statistics and the input topology [[[Europe, Asia], Ancient], Outgroup]. A negative D-score suggests a closer relationship of an ancient individual to western European individuals, whereas a positive D-score suggests a closer relationship to the Asian individual. Dot colours show the Asian individuals’ geography (longitude) as seen in the map insert.. Bar colours next to their names represent the geographic location (longitude) of the ancient individuals based on the colours in Fig 1A.

## Discussion

Through the inclusion of genomic data from Late Pleistocene and Holocene brown bear individuals sampled outside the current distribution range of the species (Fig 1A), we uncover ancient diversity not present in the available contemporary data. In general, the ancient individuals do not follow the clear phylogeographic subdivision found in contemporary brown bear individuals, with geographically structured populations most likely driven by isolation-by-distance (Fig 1). Therefore, although brown bears successfully made it through the Late Quaternary megafauna extinction event, populations and diversity were lost along with large areas of their past distribution.

A pattern of population continuity since the Late Pleistocene is not obvious in Europe, where the ancient European individuals were widely distributed within our PCA analysis (Fig 1B,C) and not clustering nearest to their closest contemporary geographic counterparts. Most notably, the ancient European (Irish and Austrian) individuals did not conform with the hypothesis of the isolation-by-distance, derived from patterns of diversity and substructuring in contemporary brown bear populations (Fig 1D). Our findings suggest brown bears previously comprised disjunct populations. The two Holocene Irish individuals (aged 5-3.8 ky BP) appear to be part of the European cluster; they were both closer to European bears in the PCA relative to their Late Pleistocene counterparts (aged 13.7-12.2 ky BP). However, the low levels of data and filtering criteria may make individuals appear more closely related to their closest cluster than they are. When considering the pairwise distance, ancestry, and D-statistics analyses, the Holocene individuals only have slightly more European ancestry. The phylogeographic structure of contemporary brown bears in Europe is considered consistent with an “expansion/contraction” model of postglacial recolonisation, where populations contracted into southern Europe during glacial periods and recolonised the rest of Europe after the glacial period [12]. Therefore, the increased contemporary European ancestry in the two Holocene Irish individuals may be due to genetic exchange with the encroaching ancestors of the contemporary European populations. Lost diversity is in line with previous findings of higher levels of mitochondrial diversity in Late Pleistocene European brown bears relative to contemporary European bears [15].

In Late Pleistocene Europe, brown bears had higher δ^15^N values, indicating more carnivorous diets [22]. However, lower δ^15^N values towards the end of the Late Pleistocene and into the Holocene have been proposed as signs of dietary flexibility as the environment changed, and brown bears shifted towards increasing herbivory [15]. Based on our findings, we suggest the decline in δ15N may reflect a loss of the more carnivorous bears, which carried their own unique genetic signal rather than the same bears shifting their diet. Although available δ15N for the ancient Irish individuals do not show a clear difference between time periods (Late Pleistocene; 6.53, 6.72; Holocene: 4.57, 8.83) [48], limited sample size prevents us from exploring this hypothesis further.

In the PCAs, all Late Pleistocene European bears show a close relationship to the Himalayan bear (Fig 1B,C), suggesting the Himalayan bear population may have been much more widespread in the past. However, this finding may be due to an overrepresentation of ancestrally shared alleles, and underrepresentation of population specific alleles; we only have data from one Himalayan individual and filter rare variants (minimum minor allele frequency 0.05 or minimum minor allele count 2) to remove the noise caused by low coverage in our ancient samples. Admixfrog does not utilise the same filtering of rare alleles, and provides support for a filtering-driven bias; we find the Late Pleistocene individuals have relatively high levels of ancestral alleles and low levels of Himalayan-specific ancestry. Not belonging to the same population as the Himalayan bear is further reinforced by the D-statistics results that show a closer relationship of the Late Pleistocene European bears to contemporary European individuals rather than to the Himalayan individual.

The Late Pleistocene Siberian individuals CGG_1_020006 (eastern Siberia) and Denisova_LP (central Siberia) also do not clearly cluster with any contemporary individuals in the PCA (Fig 1B,C). This suggests some lost diversity in Siberia, in accordance with our observations in Europe, and. it is in agreement with previous work showing mitochondrial lineages in Late Pleistocene Siberian individuals that have since been lost in contemporary individuals [50]. However, Denisova_LP does conform to the isolation-by-distance hypothesis when comparing PC1 and longitude (Fig 1D). This is also supported by the D-statistics results, which place Denisova_LP as intermediate between contemporary European and Asian individuals (Fig 3), as would be expected based on its intermediate geographic locality.. The unique placement of CGG_1_020006 is in line with a previous study investigating its mitochondrial genome, which found it belonged to a haplogroup not found in living brown bears [22].

In Japan, all contemporary Hokkaido individuals cluster together in the PCAs and appear to be one population (Fig 1B,C). This in contrast with previous hypotheses of multiple migration events to Hokkaido based on mitochondrial genomes [14,51]he timing of divergence suggested brown bears migrated from mainland Eurasia to Honshu, Japan’s largest island, at least twice [14], and to Hokkaido at least three times [51]. In our analyses, the Late Pleistocene individual from Honshu (JBB-32K) falls within or very close to the contemporary Hokkaido individuals (Fig 1B,C). The age of JBB-32K of 32,464 ± 255 cal. BP suggests population continuity since at least this time. The mitochondrial genome is limited in resolution, as it only represents a single loci and may be biased by gene flow and incomplete lineage sorting. As gene flow has been reported in Japanese brown bears [30] and both female and male brown bears are known to disperse [52], these hypothesised migration events may represent gene flow events into an already established Japanese population, mediated by a few individuals, and not large population movements. A proposed land bridge in the Tsugaru Strait connected the islands of Hokkaido and Honshu around 140,000 years ago during the glacial period of MIS6. At this time, sea levels were at their lowest and would have facilitated connectivity between the two islands [53]. This relatively recent connectivity likely led to the shared recent ancestry between Honshu and Hokkaido bears we see in our results.

In contemporary North American brown bears, three genetically divergent mitochondrial lineages have been described, leading to the hypothesis of three independent colonisation events from Eurasia [12]. In contrast, the nuclear data are consistent with one single metapopulation, with several subpopulations likely driven by isolation-by-distance (Fig 1B). The ancient brown bear from Quebec, aged 11,279 ± 30 cal. BP, may represent lost diversity, as it does not cluster closely with any of the contemporary North American bears and brown bears are thought to have gone extinct in the region by the late 20th century [54]. The Quebec individual may have come from the most recent replacement/migrations towards the end of the LGM. Previous studies using specimens dated to before this have uncovered extinct and divergent mitochondrial lineages in Eastern Beringia, suggesting dispersals into North America at widely different points in time [16].

The utilisation of published palaeogenomic data for several European and one North American brown bear enabled the recovery of lost diversity in Europe and North America. However, by generating palaeogenomic data from an additional two Asian individuals, we were able to geographically expand our inferences across their Holarctic range. In doing so, we found that although Eurasian brown bears currently display patterns of isolation by distance across the two continents, this was not always the case. Taken together, our study shows that even with ultra-low coverage palaegenomes, important inferences can be made about how a species has changed through time which is not captured in the genomes of modern individuals alone.

## Data accessibility

The raw Illumina sequence data for individual JBB-32K are available under DDBJ DRA ID: DRA015631. The raw Illumina sequence data for individual CGG_1_020006 are available under Bioproject PRJNA930648 on NCBI.

## Funding

This study was supported by Grants-in-Aid for Scientific Research (no 22K18730 and 23KK0062) from the Japan Society for the Promotion of Science (JSPS), the Villum Fonden Young Investigator Programme, grant no. 37352.

## Acknowledgements

We thank Dr. Naoki Kohno for his valuable discussions regarding sea level changes and periods of low sea levels in Japan.

## Notes

### Competing Interest Statement

The authors have declared no competing interest.

### Summary of Updates

This version has been massively revised by including ∼100 new genomes over the Holarctic distribution and with more focus on the ancient samples

