## Supplementary information for "The origins and diversification of Holarctic brown bear populations inferred from genomes of past and present populations"

**Supplementary methods**

**Error estimates**

We estimated levels of errors in our ancient dataset based on a high quality "error-free" individual using ANGSD (-doAncerror). For this we generated a consensus fasta file for the Apennine individual (APP2) using ANGSD and the filters: -minq 25 -minmapq 25. We used the spectacled bear as the outgroup. When running the error estimation method we used no filtering parameters. This method works under the assumption that the error-free individual and the sample of interest are equally distant to the outgroup. It counts the excess of derived alleles in the sample of interest compared to the derived alleles of an error-free individual. All individuals showed relatively similar levels of errors, with elevated values associated with ancient DNA damage, G-A and C-T transitions (Supplementary Fig S4). Individual Kunashir2 showed highly elevated relative error levels of all types of substitutions, not just those associated with ancient DNA damage, relative to the outgroup and high coverage APP2 indicating high levels of sequencing errors. We therefore removed this individual from further analyses.

**Supplementary tables**

**Supplementary table S1:** Information on the genomic data used in this study (attached as spreadsheet).

**Supplementary table S2:** Reduced dataset of individuals used in the D-statistics topology test.

| **H1 individuals** | **H2 individuals** |
|---|---|
| SouthNorway7 | TurkeyMartin |
| ALP1 | Ural6 |
| GRE1 | Ge |
| SLK1 | IranGudrun |
| SPA1 | CentralRussia5 |
| 191Y | Himalaya1 |
| Ursus_arctos_Sweden_Dalarna | CentralRussia1 |
| BGI-brownbear-20105373 | S235 |
| Norway8 | CentralRussia4 |
| Rumania5 | FarEast9 |
| Estonia1 | FarEast6 |
| Finland3 | Amur3 |
| Kirkenes | Amur1 |
| Russia_Kola1 | DRR276774 |
| | Amur4 |
| | Hokkaido1 |
| | DRR276779 |
| | FarEast1 |
| | Kamtschatka4 |

**Supplementary figures**

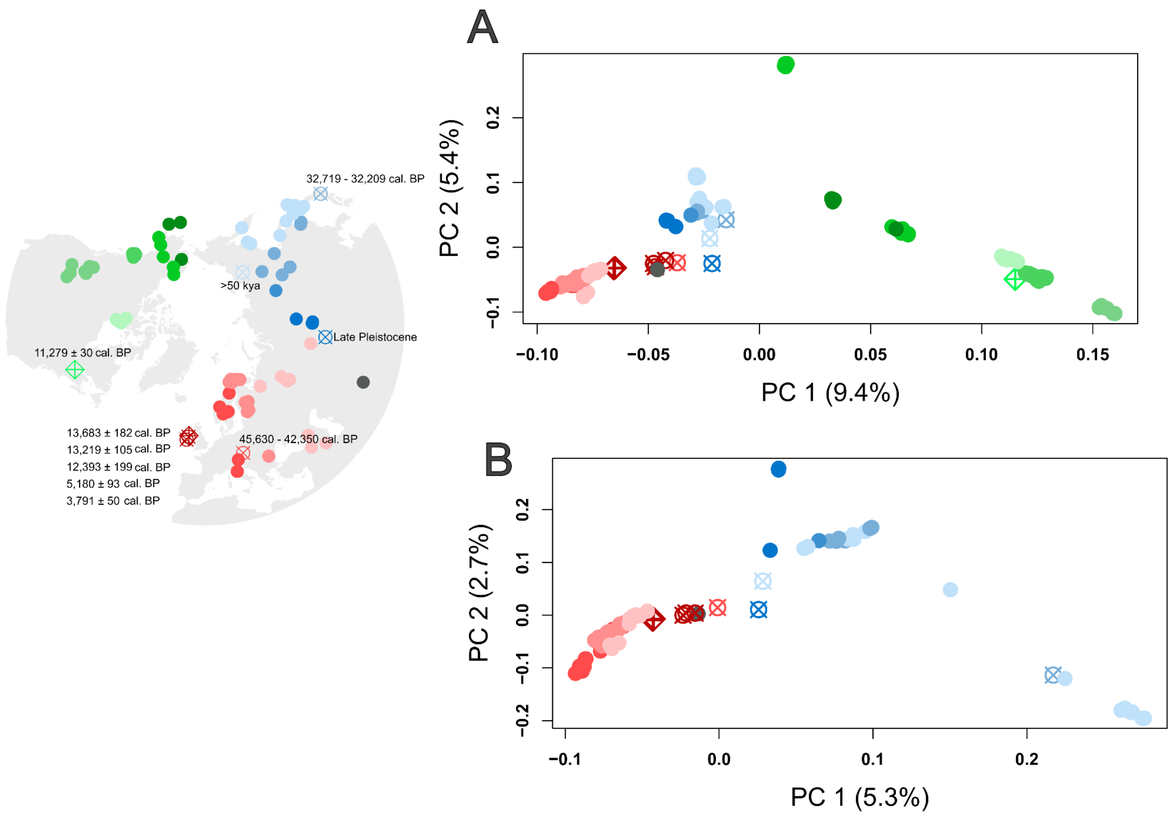


**Supplementary figure S1: A)** PCA generated from EMU on full dataset, **B)** PCA generated from EMU on Eurasia only dataset. Percentage values in parentheses on the X and Y axes show the percentage of variation explained by the principal component. Colours and shapes represent geography and age respectively as shown on the map to the left.

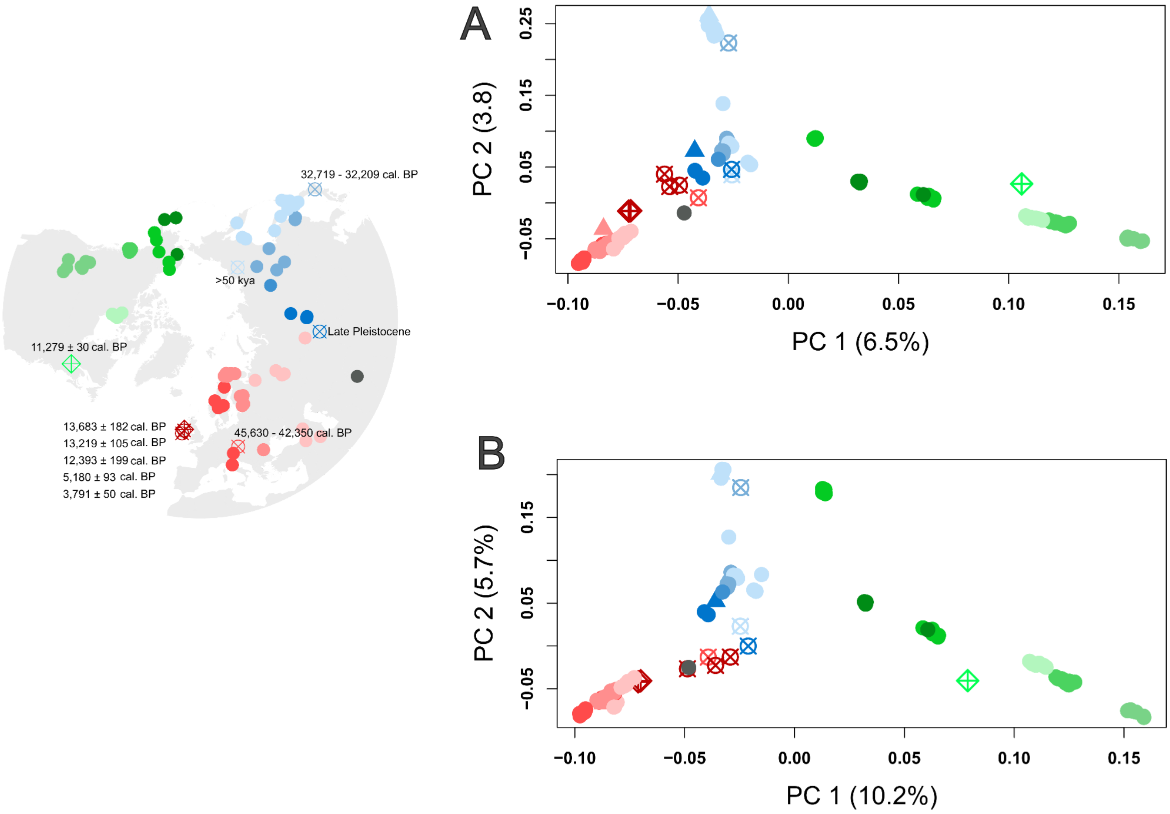


**Supplementary figure S2: A)** PCA generated from pseudo-haploid base calls, **B)** GL. Percentage values in parentheses on the X and Y axes show the percentage of variation explained by the principal component. Colours and shapes represent geography and age respectively as shown on the map to the left. Individuals replaced by simulated damage and low coverage indicated by triangles Kirkenes (Europe), Rus235 (Central Asia), DRR276778 (Japan).

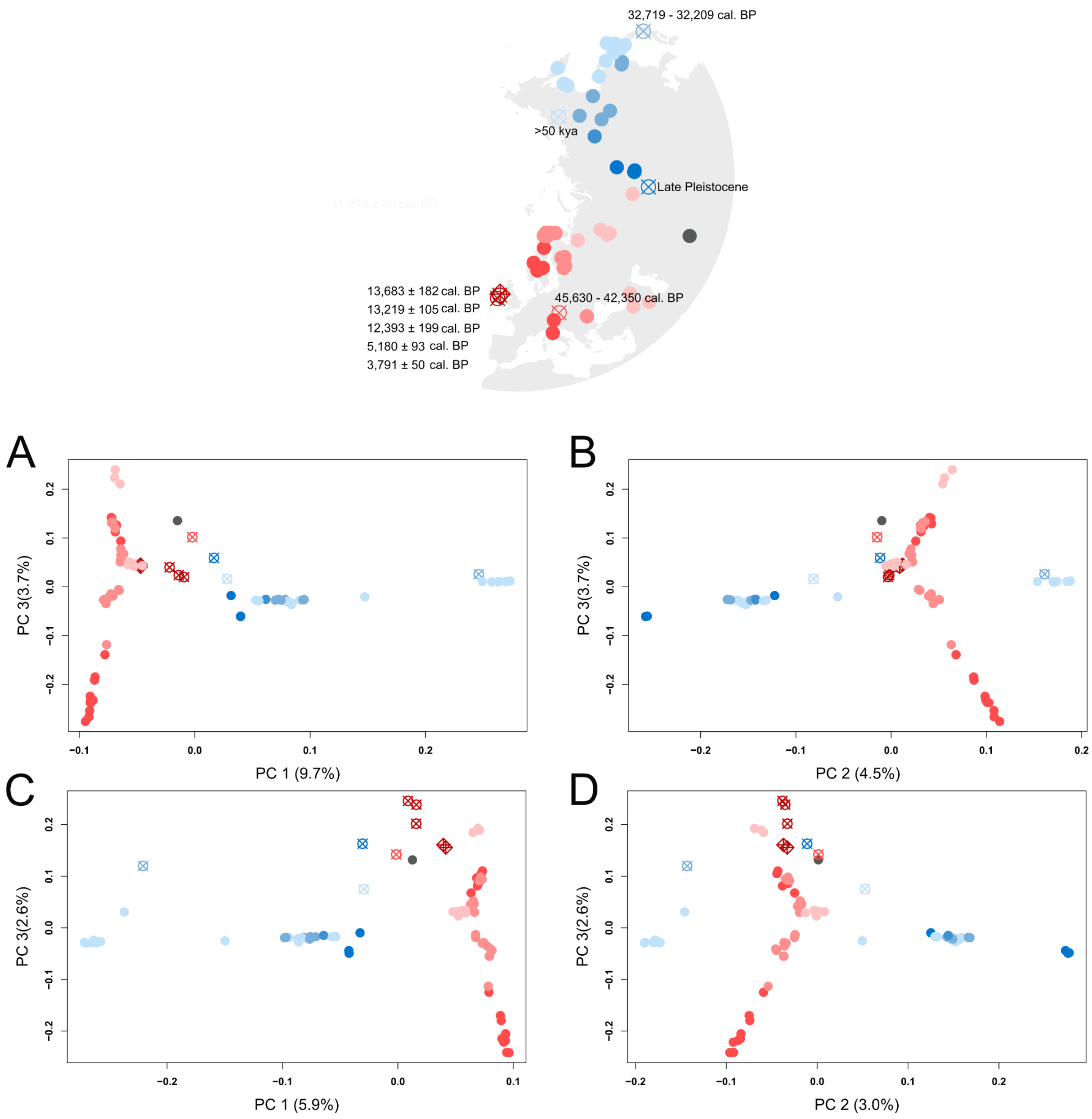


**Supplementary figure S3:** PCA showing either PC1 vs. PC3 or PC2 vs. PC3 using the Eurasia only dataset. **A and B)** PCAs generated from GL, **C and D)** pseudo-haploid base calls. Percentage values in parentheses on the X and Y axes show the percentage of variation explained by the principal component. Colours and shapes represent geography and age respectively as shown on the map to the left.

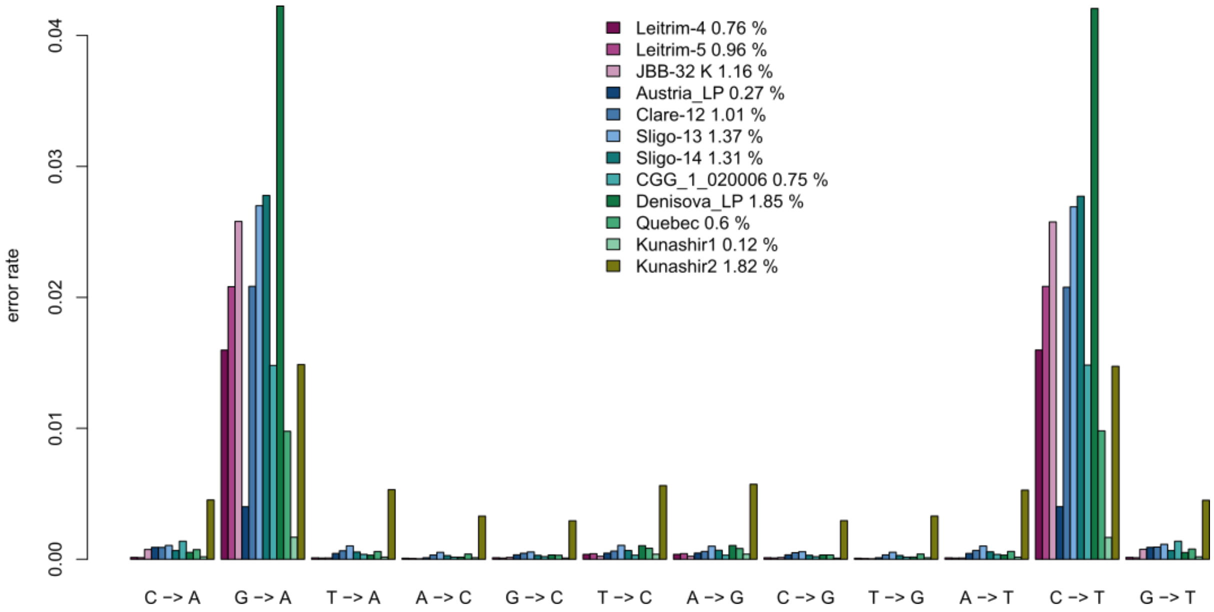


**Supplementary figure S4:** Relative error rates of each ancient individual estimated using an outgroup and an error-free high quality individual (APP2) in ANGSD. X axis shows the type of substitution.
